# ZONAB regulates DNA methylation and mitochondrial function to control endothelial cell senescence

**DOI:** 10.1101/2025.11.20.689552

**Authors:** Wenyi Jiang, Eleanor Lynam, Juliette Delafosse, Graeme M. Birdsey, Anna M. Randi, Karl Matter, Maria S. Balda

## Abstract

Regulation of the endothelial stress response is important for blood vessel homeostasis and angiogenesis, processes disrupted in common vascular diseases and ageing. Here, we discovered that the Y-box factor ZONAB (ZO-1-associated nucleic acid binding protein; *YBX3*), a gene associated with risk loci for severe vascular disorders, regulates endothelial homeostasis and angiogenesis. By combining cell-based assays with primary endothelial cells and genome-wide expression and methylation measurements, we found that ZONAB depletion results in mitochondrial deregulation, increased reactive oxygen species and a defective oxidative stress response, as well as increased promoter methylation of cell cycle genes. Consequently, ZONAB depletion triggered cellular senescence via a phosphatidylinositol 3-kinase (PI3K)/Akt-dependent pathway, which was attenuated by an antioxidant or by drugs targeting mitochondrial function or fragmentation. Thus, our results reveal how ZONAB controls changes in gene expression and DNA methylation to regulate endothelial proliferation, inflammation, and angiogenesis, indicating a central role of ZONAB in vascular health.

## Introduction

Endothelial cells line the internal walls of blood and lymphatic vessels. They can exhibit high plasticity and can switch between a quiescent state, which supports vessel stability and function, and a proliferative and migratory state required for angiogenesis. Both angiogenesis and vessel homeostasis are affected by ageing, which generally results in a decrease in angiogenic potential and an increase in rates of diseases induced by endothelial malfunction, such as atherosclerosis and microvascular conditions affecting the heart and/or the brain ^1,2^. Endothelial senescence induced by ageing and chronic stress is a key driver of such diseases, indicating that mechanisms that support the endothelial stress response are crucial for preventing the induction of stress-induced senescence. However, these mechanisms are only partially understood.

Cellular senescence is a state of permanent stop in cell proliferation induced by ageing or stress. It may be stimulated by telomere shortening, loss of telomere functionality, or by stress conditions, such as oxidative or oncogenic stress ^2,3^. Accumulation of senescent cells often induces an inflammatory environment due to the release of proinflammatory stimuli such as IL1α/β and IL6 by the senescent cells, a process called the senescence-associated secretory phenotype (SASP). While SASP promotes processes such as wound healing, the accumulation of senescent cells with a SASP in ageing stimulates chronic inflammatory conditions due to the continuous release of proinflammatory factors.

Given the importance of stress for the induction of senescence, one would expect that genetic variants of stress response genes exist that predispose people to develop diseases such as atherosclerosis. Genome-wide association studies (GWAS) focusing on atherosclerosis identified such genes based on their association with loci that increase the risk of developing the disease ^4,5^. Similarly, another study focused on genetic variants predisposing people to high systolic blood pressure, also a process affected by endothelial homeostasis ^6^. A common gene identified by those studies is *YBX3*. Variants in this gene have also been linked to ovarian ageing ^7^, suggesting that its function could be linked to tissue ageing and cellular senescence.

*YBX3* encodes a multifunctional nucleic acid-binding protein that is also called ZONAB (ZO-1-associated nucleic acid binding protein) as it can interact with the tight junction protein ZO-1 ^8,9^. In epithelial cells, ZONAB controls cell density, proliferation, and the cellular stress response ^9–13^. ZONAB controls G1/S phase transition by transcriptional regulation of cyclin D1 and PCNA, and by forming a complex with CDK4 ^9,11^. ZONAB is not only regulated by binding to ZO-1 but is also inhibited by Ral A, a Ras-effector pathway, and activated by the RhoA activator GEF-H1 ^10,13,14^. Overexpression of ZONAB has been linked to cancer cell proliferation and metastasis in multiple tissues. However, the underlying mechanisms for its role in tumorigenesis are not well defined ^15–19^. These studies have generally been performed in immortalised epithelial cell lines; therefore, whether loss of ZONAB function is linked to cellular senescence and if it is functionally relevant in endothelia remains unknown.

Here, we show that ZONAB is required for angiogenesis *in vivo* and normal angiogenic properties *in vitro*. ZONAB depletion in human primary endothelial cells induced cellular senescence. Cellular assays and genome-wide expression and methylation analyses indicate that ZONAB depletion resulted in genome-wide changes in gene expression, at least partially mediated by methylation-driven epigenetic changes, leading to mitochondrial dysfunction, a disrupted oxidative stress response, and cellular senescence.

## Results

### Depletion of ZONAB inhibits endothelial cell migration and angiogenesis

We first asked whether ZONAB is functionally relevant for endothelial angiogenesis. We established a loss-of-function model for ZONAB using human dermal microvascular endothelial cells (HDMECs). HDMECs form well-assembled and functional tight and adherens junctions ^20^. We transfected the cells with control non-targeting or two ZONAB-specific siRNAs (termed 3 and 4). Immunoblotting confirmed efficient depletion of both ZONAB isoforms (ZONAB-A and ZONAB-B) by individual siRNAs as well as a pool of both ZONAB siRNAs (Fig. 1A).

**Figure 1.**
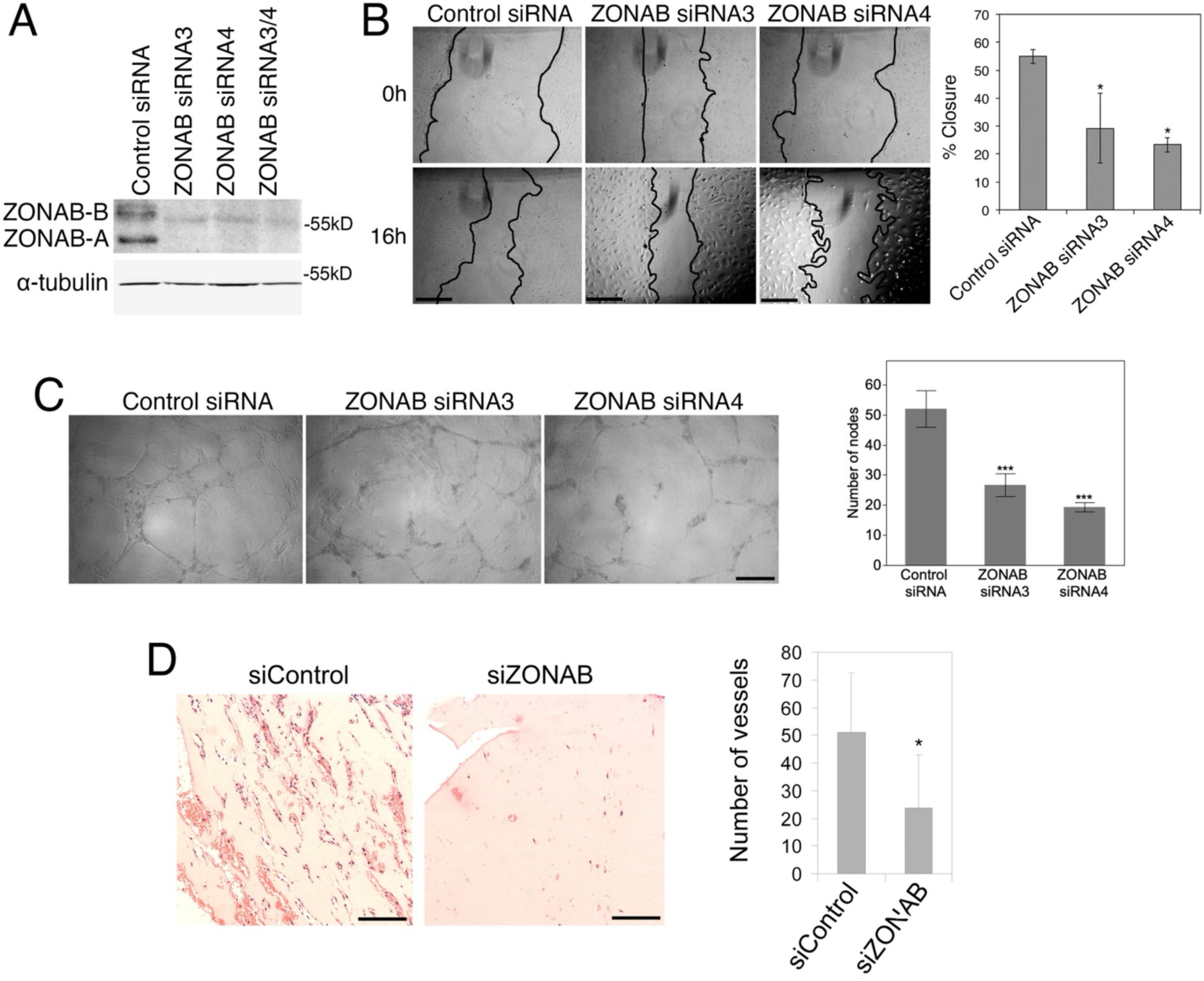
ZONAB regulates endothelial cell migration and angiogenesis. (A) Analysis of ZONAB expression by immunoblotting of cell lysates of HDMECs transfected with non-targeting control siRNAs or with two different ZONAB-targeting siRNAs (ZONAB siRNA 3 or 4) or as a pool (ZONAB siRNA 3/4). (B) HDMECs were transfected with the indicated siRNAs. Scratch wounds were then inflicted to induce cell migration 48 hours later. Representative images of wounds are shown, taken at 0 and 16 h after scratching. Means ± 1 SD, n=3. (C) HDMECs were transfected with the indicated siRNAs and, 48 hours later, replated on 100% Matrigel to test their angiogenic potential in a tube formation assay on Matrigel. Network formation was analyzed after 16 hours. The number of branches in each sample was quantified. Means ± 1 SD, n=3. (D) The effect of ZONAB depletion on angiogenesis *in vivo* was measured by injecting mice with Matrigel-containing FGF and siRNAs as indicated to induce angiogenesis. After 7 days, the plugs were harvested and analyzed. Means ± 1 SD; control siRNA, n=6; ZONAB siRNA, n=12). Bars, 250 μm. T-tests: *p<0.05, ***p<0.001

We first tested whether ZONAB depletion impacts cell migration by performing a scratch wound assay with siRNA-transfected cells. Figure 1B shows that ZONAB-depleted endothelial cells migrated more slowly than control siRNA-treated cells. To evaluate the endothelial angiogenic potential *in vitro*, we next performed Matrigel tubulogenesis assays. ZONAB depletion reduced network formation and led to a ∼50% decrease in node formation (Fig. 1C). ZONAB is thus functionally important for endothelial cell migration and tube formation on Matrigel *in vitro*.

We next asked whether ZONAB also regulates the endothelial angiogenic potential *in vivo*. We performed Matrigel plug assays in mice combined with siRNAs as we did previously for ZO-1 ^20^. ZONAB siRNAs led to a more than 50% reduction in vessel formation when ZONAB siRNAs were co-injected with FGF and Matrigel, indicating that vessels failed to invade the FGF-rich matrix *in vivo* (Fig. 1D). ZONAB is thus essential for angiogenesis *in vitro* and *in vivo*.

### ZONAB regulates the actin cytoskeleton

The actomyosin cytoskeleton is crucial for cell migration and angiogenesis. Hence, we tested whether ZONAB depletion affects the actomyosin cytoskeleton and junction formation. ZONAB-depleted cells formed striking thick F-actin bundles along the cell periphery instead of the normal junctional actin pattern (Fig. 2A). These peripheral F-actin bundles also contained phosphorylated MLC (ppMLC). Myosin IIA was also enhanced along those F-actin fibres and at tricellular corners, whereas AKAP-12 still outlined the lateral membrane (Fig. 2A). Thus, ZONAB depletion induced actomyosin reorganisation. The cells generally appeared more spread, reflecting the changes in F-actin organisation and myosin activation. ZONAB depletion did not affect ZO-1 localisation but led to reduced junctional staining of other proteins that are at least partially associated with tight junctions. Staining for the scaffolding protein JACOP/paracingulin/CGNL1, the tight junction transmembrane protein claudin-5, and the regulator of tight junction assembly IQGAP1 were strongly reduced (Fig. 2B). In contrast, core adherens junction proteins such as VE-cadherin, β-catenin, and p120 catenin appeared normally localised (Fig. 2C). The exception was vinculin, which moved to focal adhesions along the cell periphery, reflecting the increase in myosin activation along the induced peripheral F-actin bundles and suggesting increased tensile forces acting on focal adhesions.

**Figure 2.**
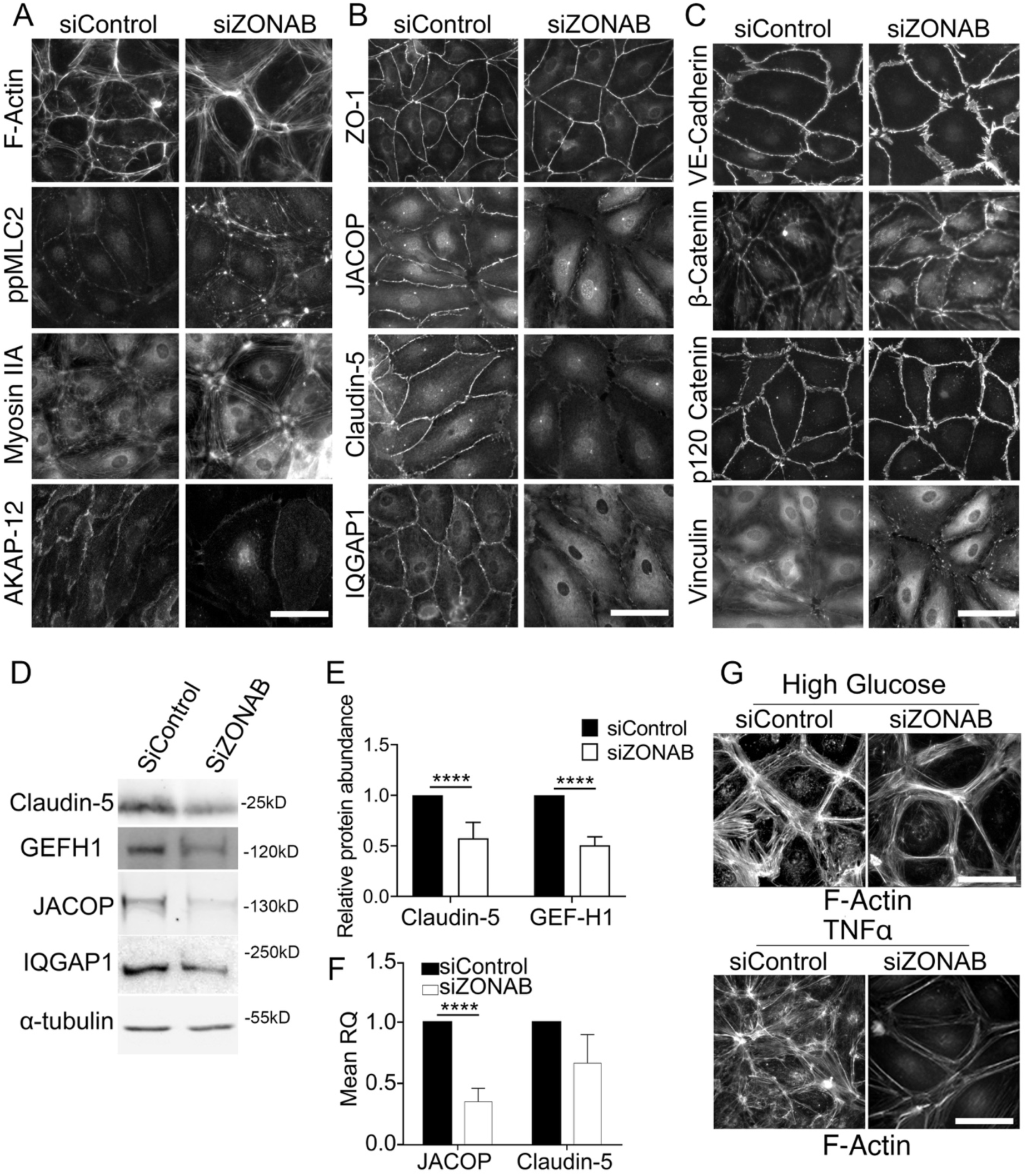
ZONAB depletion affects the actomyosin cytoskeleton and junctional complexes. (A-C) Immunofluorescence images of HDMECs treated with control or ZONAB siRNAs for 72 hours before immunofluorescent staining with phalloidin for the visualization of F-actin or antibodies against the indicated proteins. Scale bar, 50 μm. (D-E) Immunoblotting of lysates from cells transfected with siRNAs for 72 hours. α-tubulin (bottom) was used as a loading control for the densitometric analysis shown in panel E. F) mRNA expression analysis by RT-qPCR for JACOP and Claudin-5 in control or ZONAB siRNA-treated cells. Values were normalized using GAPDH expression, and obtained ratios were normalized to control siRNA values (means ± 1 SD, n=3). G) F-actin staining of HDMECs transfected with control or ZONAB siRNA in medium with high glucose (33mM, for 24 hours) or stimulated with 5 ng/ml TNFα for 5 hours. Scale bar: 50 μm. All graphs show means ± 1 SD (n=3).T-tests: **** P ≤ 0.0001

Protein expression of JACOP, claudin-5 and IQGAP1 and GEF-H1, a Rho guanine nucleotide exchange factor known to interact with JACOP ^20^, was reduced as determined by immunoblotting, in agreement with the immunofluorescent data (Fig. 2D and E). JACOP was already reduced at the mRNA level, but claudin-5 mRNA expression was not significantly affected (Fig. 2F). Overall, these data indicate that ZONAB depletion led not only to a reorganisation of the actomyosin cytoskeleton but also to disruption of junctional proteins known to interact and/or regulate actin dynamics. Given the increase in peripheral actomyosin fibres and the vinculin redistribution to focal adhesions, it is likely that cells adhere more tightly to the substrate, inducing the observed increase in cell spreading and reduction in angiogenesis.

To assess the stability of the ZONAB-induced F-actin reorganisation, we treated cells with stimuli known to promote F-actin remodelling. Figure 2G shows that high glucose (25mM, 24h) or TNFα (5 ng/ml for 5h) led to induction of differently arranged stress fibres in endothelial cells compared to controls cells (see panel A); however, the high glucose treatment of control cells was already similar to ZONAB-depleted cells in normal medium and did not further change in high glucose medium. The TNFα-induced striking F-actin reorganisation in control cells did not occur in ZONAB-depleted cells, but TNFα appeared to reduce some of the peripheral ZONAB depletion-induced F-actin bundles, indicating that TNFα-induced actin remodelling requires ZONAB expression.

### ZONAB depletion reduces endothelial cell proliferation

The cell shape changes with increased cell spreading suggested that ZONAB may regulate endothelial cell proliferation, resulting in monolayers with a lower cell density (Fig.2A-C, Fig.3A). Quantification of cell numbers after 3 days of ZONAB depletion when cells had reached confluence by CyQUANT assay revealed that cell numbers were indeed reduced by about 25% (Fig.3B). No increases in apoptosis (Apo-ONE) or necrosis (Cyto-TOX) were detected, indicating that the cell number changes were not due to increased cell death (Fig. 3C).

**Figure 3.**
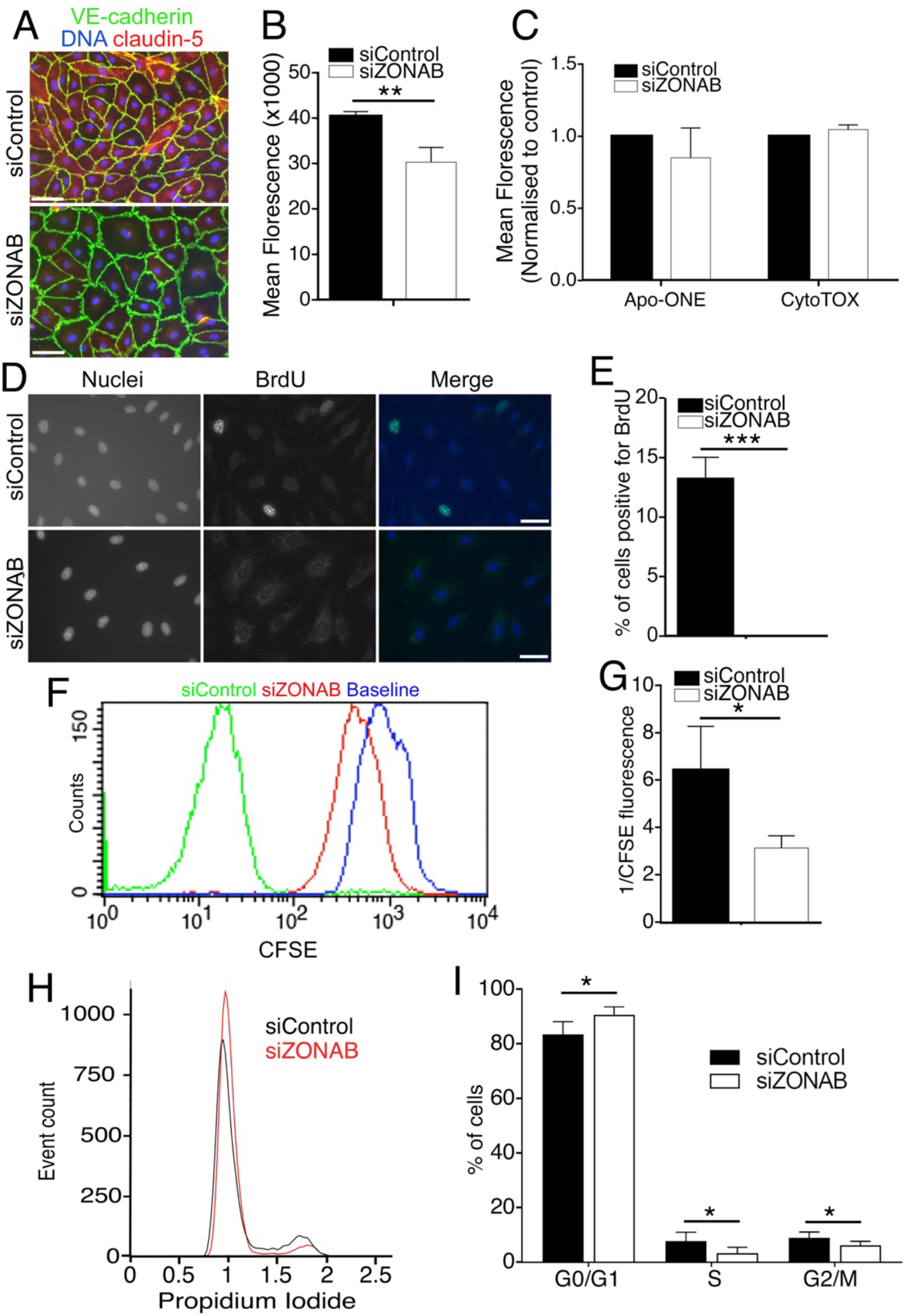
ZONAB regulates endothelial cell proliferation. (A) Immunofluorescence of siRNA-transfected HDMECs stained for VE-cadherin, claudin-5 and nuclei. Scale bar, 50 μm. (B) Cell density measured by measuring DNA content using the CyQUANT Assay. (C) Caspase-3/7 activity (Apo-ONE) and LDH release (CytoTox) measurements of siRNA-transfected cells. Values were normalized to control siRNA determinations (D-E) Assessment of proliferation by BrdU incorporation. Shown are images of nuclear and BrdU staining. Scale bar: 50 μm. (F-G) Cell division capacity assessed by CFSE CellTrace tracking dye. (H) Propidium iodide flow cytometric analysis of cell cycle stages. (I) The quantification shows the percentage of cells in G0/G1, S and M phases. All graphs show means ± 1 SD (n=3). T-tests: * P ≤ 0.05, ** P ≤ 0.01, *** P ≤ 0.001

Next, we measured proliferation by adding bromodeoxyuridine (BrdU) for 4 hours during the last day of the experiment to label cells in S-phase. ZONAB-depleted cells exhibited a strong decrease in the number of cells that incorporated BrdU (Fig. 3D and E), indicating that ZONAB-depleted cells no longer entered S phase on the last day of incubation. We next measured cell division using a carboxyfluorescein succinimidyl ester (CFSE) based cell-tracking dye that is incorporated into DNA and split equally between the two daughter cells upon mitosis, resulting in weaker signals as the cells divide. Control cells had a significantly weaker CFSE fluorescence than ZONAB-depleted cells, indicating that ZONAB-depleted cells had replicated less often than control cells (Fig. 3F and G). Thus, ZONAB depletion reduced endothelial cell numbers and proliferation. Propidium iodide (PI) staining combined with flow cytometry indicated that ZONAB-depleted cells were largely arrested in the G0/G1 phase of the cell cycle (Fig. 3H and I). Thus, ZONAB depletion arrested endothelial cell proliferation by halting the cell cycle at G0/G1 phase.

### ZONAB regulates the expression of central cell cycle regulators

We next employed an Affymetrix cDNA gene expression microarray and RNA sequencing to determine the effect of ZONA depletion on gene expression in endothelial cells (Tables S1 and S2). The two independent approaches largely confirmed each other. ZONAB depletion resulted in the identification of a large set of deregulated genes encoding cell cycle regulators such as cyclins and cell cycle kinases (CDKs), CDK inhibitors, cell cycle-regulating transcription factors, and genes required for DNA packing, centromere assembly and chromosome separation (Fig.4A, Tables S3-S6; KEGG cell cycle pathway coverage of 41% with a log_2_ cutoff of |0.5|). Immunoblotting and/or RT-qPCR confirmed downregulation of multiple cyclins, CDKs and PLK-1, as well as upregulation of CDK inhibitors (Fig. 4B-E). Hence, ZONAB depletion led to genome-wide changes in the expression of genes that regulate cell cycle progression.

**Figure 4.**
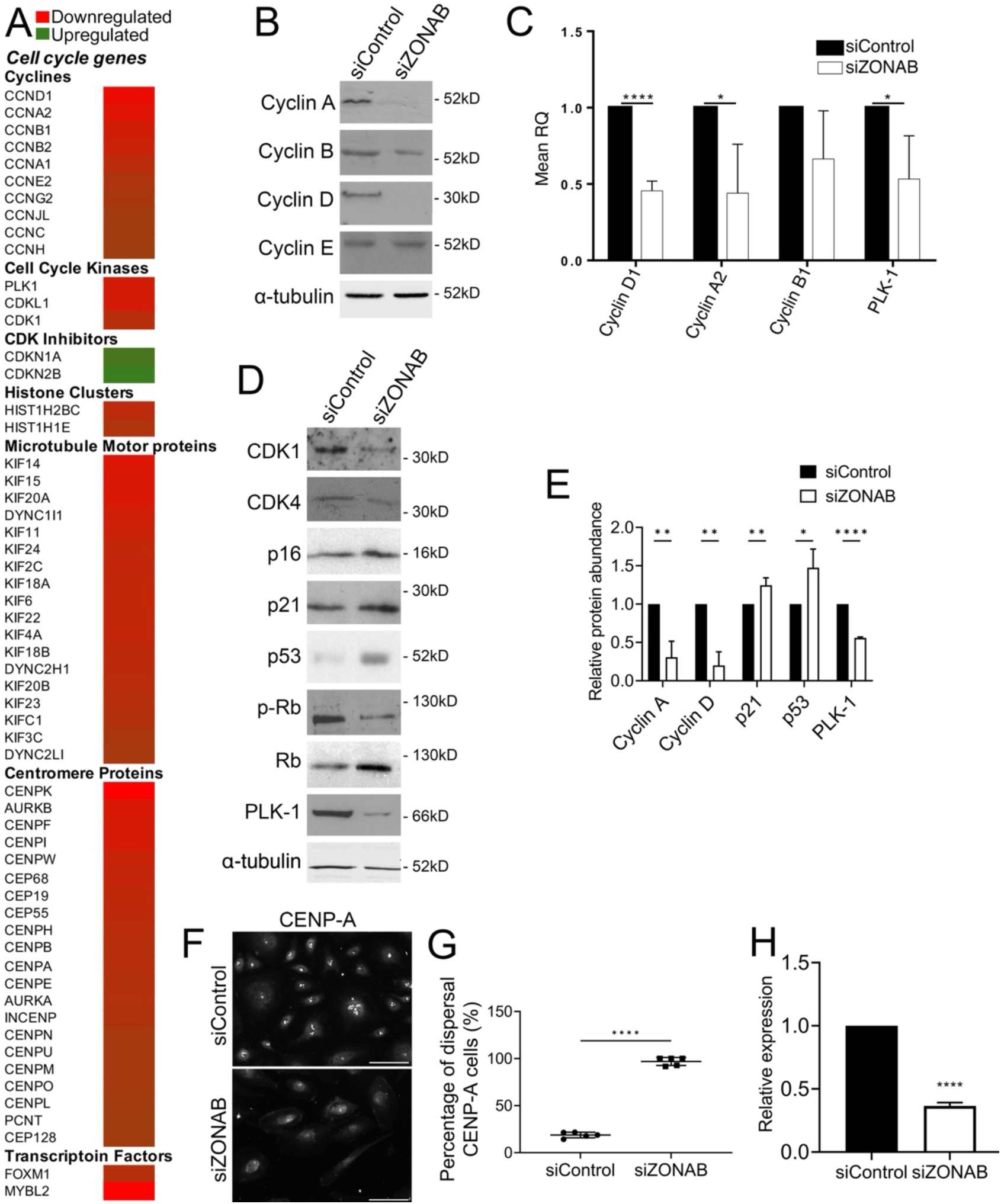
ZONAB depletion reduces the expression of cell cycle genes. (A) Expression analysis by RNA sequencing of control and ZONAB siRNA-transfected cells. Analyzed were three mRNA isolations for each siRNA set. Shown are genes repressed or induced by ZONAB depletion (change in expression log_2_> |0.5|; see also Tables S2 and S7). (B, D, E) Expression of the indicated cell cycle regulators was analyzed by immunoblotting. α-tubulin was used as a loading control for the quantifications shown in panel E (means ± 1 SD, n=3). (C) RT-qPCR for mRNA quantification of selected cell cycle regulators. Values were normalized using GAPDH expression, and obtained ratios were normalized to control siRNA values (means ± 1 SD, n=3). (F-G) Immunofluorescent staining for CENP-A. Scale bar: 100μm, the quantification shows percentage of cells with dispersed staining for CENP-A (means ± 1 SD, n=5 images from 3 independent experiments). (H) mRNA expression of CENP-A in control or ZONAB siRNA-transfected cells (means ± 1 SD, n=3). T-tests: *p<0.05, **p<0.01, ***p<0.001, ****p<0.001

Given the inhibition of cell cycle progression observed upon ZONAB depletion (Fig. 3I), we next tested expression and phosphorylation of the tumour suppressor retinoblastoma protein (Rb), which regulates G1/S phase transition. Rb protein expression was increased after ZONAB deletion, but phosphorylated Rb was decreased (Fig. 4D). Thus, ZONAB is required for efficient Rb phosphorylation and, thereby, inactivation. Similarly, expression of the tumour suppressor p53 was increased in ZONAB-depleted cells as well as expression of the cell cycle inhibitor p21 (Fig. 4D and E). Hence, the pattern of gene expression changes indicated an inhibition of G1/S phase transition by induction of inhibitory mechanisms, which agreed with the cell cycle behaviour observed in figure 3.

Deregulation of cell cycle genes was not restricted to G1/S phase transition but also affected important regulators of G2/M phase transition, such as PLK-1 (Fig.4B-E). Expression of genes required for centromere formation, chromosome separation and mitosis was generally reduced (Fig.4A). PLK-1 promotes CENP-A loading on centromeres ^21^. ZONAB depletion not only reduced CENP-A expression but also promoted dispersal of the remaining centromere protein (Fig.4F-H).

ZONAB depletion thus deregulated the expression of genes guiding efficient progression at different steps of the cell cycle, indicating a general reduction in the proliferation potential of endothelial cells.

### ZONAB depletion stimulates cellular senescence

The RNA sequencing data indicated not only cell cycle arrest but also altered expression of markers associated with cellular senescence (Fig.5A; Table S7). In particular, the expression changes indicated induction of a SASP and inflammation. ZONAB depletion significantly upregulated mRNA of inflammatory genes, such as IL1A, ICAM1, SELE and MMP10 (Fig. 5A,B). Depletion of ZO-1 did not or only modestly affect those inflammatory genes (Fig. 5B). Increased expression of ICAM-1 upon ZONAB depletion was confirmed by immunoblotting (Fig. 5C). Hence, ZONAB-depleted endothelial cells display characteristics of a SASP and are in an increased inflammatory state in addition to reduced proliferation.

**Figure 5.**
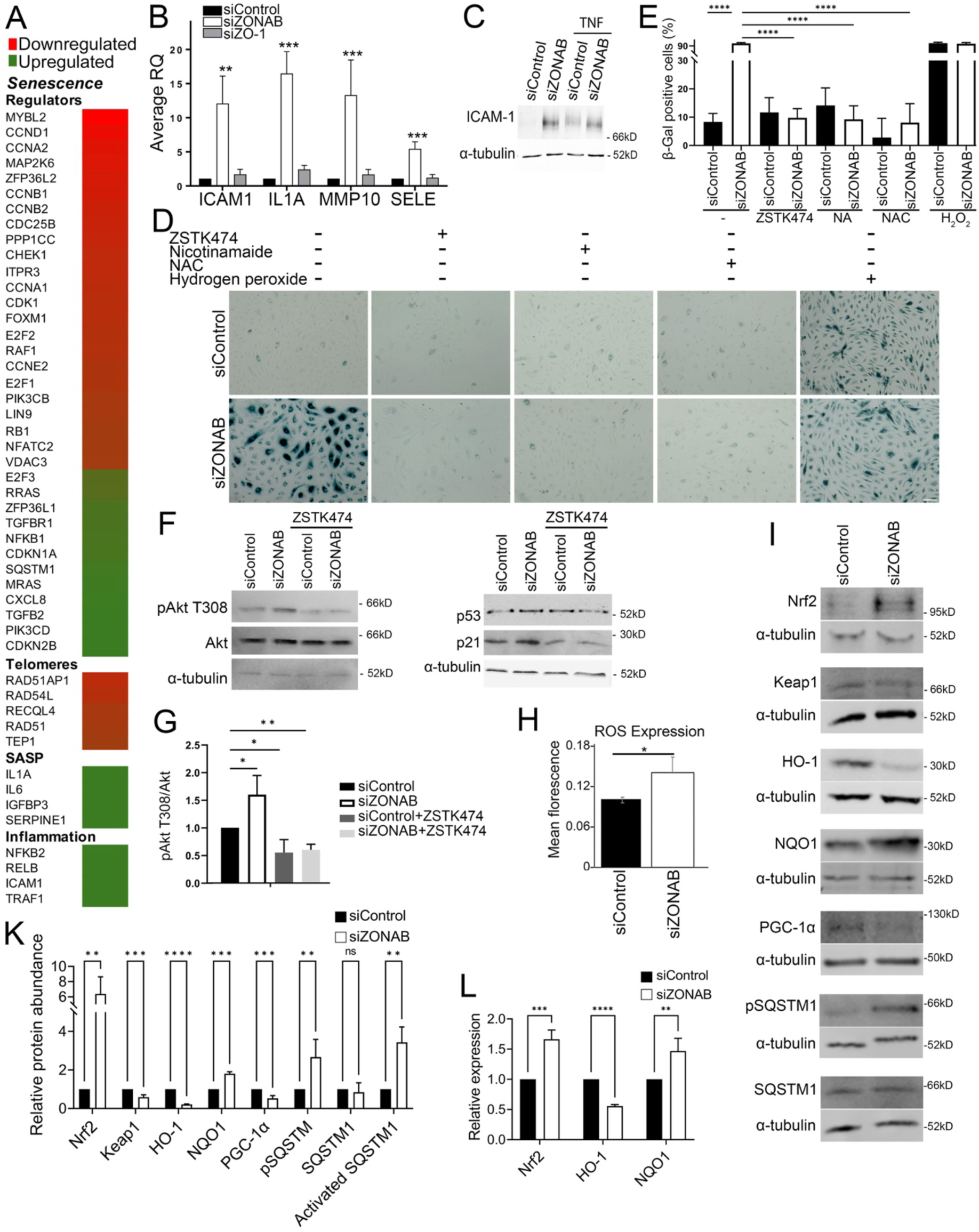
ZONAB depletion induces a senescent inflammatory phenotype that can be rescued by inhibition of PI3K. (A) Repression or induction of genes associated with cellular senescence, telomeres, the senescence-associated secretory phenotype (SASP), and endothelial inflammation (change in expression log_2_> |0.5|; see also Tables S2 and S7). (B) RT-qPCR analysis of genes associated with inflammation. Values were normalized using GAPDH expression, and obtained ratios were normalized to control siRNA values (means ± 1 SD, n=3). (C) Immunoblot of ICAM-1 expression upon ZONAB depletion without or with TNFα treatment. (D-E) Senescence-associated β-Galactosidase (SAβG) staining of siRNA-transfected cells without or with incubation for 48 hours with 1 μM PI3K inhibitor ZSTK474, 200 μM Nicotinamide, or 8 mM NAC. A positive control was incubated for 6 hours with 150 μM H2O2. Scale bar: 200 μm. Cells were imaged by brightfield microscopy, and SAβG-positive cells were counted (panel E, means ± 1 SD, n=3). (F-G) Immunoblotting for pAkt T308 and Akt, as well as for p53 and p21 shows that PI3K inhibition effectively represses Akt phosphorylation and expression of p21 and p53. The quantification in panel G shows means ± 1 SD (n=3). (H) ZONAB depletion induces ROS (means ± 1 SD, n=3). (I) Immunoblotting and (K) densitometric quantification for indicated oxidative stress response genes. α-Tubulin was used as a loading control (means ± 1 SD, n=3). (L) RT-qPCR analysis of oxidative stress response genes. Values were normalized using GAPDH expression, and obtained ratios were normalized to control siRNA values (means ± 1 SD, n=3). T-tests: *p<0.05, **p<0.01, ***p<0.001, ****p<0.001

A key marker of senescent cells is senescence-associated β-galactosidase (SAβG) activity. We tested whether ZONAB-depleted cells exhibit increased expression of SAβG using a colorimetric staining. About 80% of ZONAB-depleted cells were positive for SAβG as opposed to less than 10% of control siRNA-transfected cells (Fig. 5D,E). Hence, ZONAB-depleted cells entered cellular senescence.

Akt activation due to increased phosphatidyl inositol-3 kinase (PI3K) activity can induce cellular senescence due to p53 and p21 induction ^22^. Both proteins were also induced by ZONAB depletion (Fig. 4D,E). Akt-induced senescence is favoured by increased expression of NF1, which we also observed in ZONAB-depleted cells (Table S2) ^23^. Therefore, we tested whether Akt was activated and whether inhibition of the pathway attenuated senescence. PI3K stimulates PDK-dependent phosphorylation of Akt at T308 and can be inhibited with the small molecule inhibitor ZSTK474. In agreement, increased Akt phosphorylation at T308, as well as p53 and p21 protein expression were reduced by PI3K inhibition with ZSTK474 in ZONAB-depleted cells (Fig. 5F,G). Thus, Akt activation upon ZONAB depletion and induction of p53 and p21 was PI3K-dependent. Strikingly, the PI3K inhibitor effectively reduced the percentage of senescent cells labelled by SAβG staining (Fig. 5D,E). These data indicate that induction of cellular senescence in ZONAB-depleted endothelial cells is indeed driven by PI3K/Akt signalling.

### ZONAB depletion stimulates reactive oxygen species

Senescence can be induced by oxidative stress, which is associated with activation of the p53/p21 pathway by reactive oxygen species (ROS) ^3,24,25^. ROS content was indeed upregulated in ZONAB-depleted cells (Fig. 5H). ROS and PI3K/Akt signalling are tightly linked and regulate each other ^26^. Hence, we tested whether drugs that rescue senescence induced by H2O2, such as Nicotinamide, a form of vitamin B3 that can also help mitigate senescence by influencing NAD^+^ levels, and N-acetylcysteine (NAC), an antioxidant that scavenges ROS, can rescue the induction of cellular senescence of ZONAB-depleted endothelial cells. Both Nicotinamide and NAC strongly reduced the percentage of SAβG-positive ZONAB-depleted cells (Fig. 5D,E). ZONAB depletion-induced senescence can thus be effectively rescued by drugs known to counteract ROS.

Our data indicate that the ZONAB depletion-induced increase in ROS is functionally relevant for cells to enter senescence. Increased ROS levels are expected to stimulate increased expression of the transcription factor Nrf2, which induces an anti-ROS response ^27^. ZONAB depletion indeed stimulated increased Nrf2 expression and reduced levels of its inhibitor, Keap1 (Fig. 5I,K). However, only one of its target genes tested, NQO1, was induced, whereas HO-1 (HMOX1) was repressed (Fig. 5I-L). Thus, while ROS-induced Nrf2, induction of the key antioxidant gene HO-1 failed.

HO-1 is required to prevent endothelial senescence, and overexpression alleviates endothelial senescence via a p53-dependent mechanism ^28,29^. Hence, we further investigated possible mechanisms underlying HO-1 downregulation. Nrf2 cooperates with peroxisome proliferator–activated receptor gamma coactivator 1 alpha (PGC-1α) in the regulation of antioxidant genes ^30^. RNA sequencing indeed indicated that ZONAB depletion led to decreased expression of PGC-1α (PPARGC1A) and key target genes in addition to HO-1, such as the mitochondrial transcription factor A (TFAM) (Tables S2 and S9). Immunoblotting confirmed the downregulation of PGC-1α (Fig. 5I, K). Hence, the reduced expression of PGC-1α likely contributes to the failure of Nrf2 to induce HO-1.

To further understand the partial disruption of the Nrf2 pathway, we tested for SQSTM1/p62, which can activate Nrf2 and is itself a target gene of Nrf2 ^31^. In agreement with the downregulation of Keap1, SQSTM1 phosphorylation increased upon ZONAB depletion, indicating activation. Although SQSTM1 is an Nrf2 target gene and its mRNA expression was upregulated by RNA sequencing (Table S2), total protein remained unchanged, possibly reflecting increased autophagy and, hence, turnover of SQSTM1.

Our data thus indicate that ZONAB depletion stimulates ROS and oxidative mechanisms; however, induction is incomplete, and Nrf2/PGC-1α signalling target genes crucial for the prevention of cellular senescence, such as HO-1, are repressed, as is PGC-1α itself.

### ZONAB regulates mitochondrial fragmentation and metabolic activity

PGC-1α regulates the biosynthesis of mitochondria; hence, depletion of ZONAB may regulate the accumulation of mitochondria via PGC-1α repression ^32^. Loss of the mitochondrial network and fragmentation can lead to mitochondrial dysfunction and senescence by excessive production of ROS ^33,34^. In mammalian cells, mitochondrial mass and network formation is regulated by fusion and fission. Hence, we first asked if the mitochondrial network became fragmented upon ZONAB depletion.

Dynamin-Related Protein 1 (DRP1) is key to the regulation of mitochondrial fission ^35^. ZONAB depletion led to about twofold DRP1 upregulation (Fig. 6A,B). ZONAB depletion altered the DRP1 distribution from a more filamentous to a more dispersed and punctate staining pattern, which displayed a reduction in branch length (Fig. 6C,D). Staining with a mitochondrial membrane potential-sensitive MitoTracker and anti-TOMM20 antibodies confirmed that mitochondria in ZONAB-depleted cells became more fragmented (Fig. 6E-G). TOMM20 protein expression was reduced, indicating less mitochondrial mass (Fig. 6H,I). In addition to TFAM, several other mitochondrial genes and tricarboxylic acid cycle (TCA; Krebs cycle) components were downregulated by RNA sequencing upon ZONAB depletion, also indicating reduced mitochondrial biosynthesis (Table S8). The MitoTracker/TOMM20 ratio was increased, indicating that the remaining mitochondria were working at increased activity (Fig.6K). The reduced total mitochondrial capacity was confirmed by Seahorse mitochondrial stress test experiments that revealed that compensatory respiration (i.e., maximal activity) was almost halved by ZONAB depletion (Fig. 6L). The basal activity was not affected by ZONAB depletion (Fig. 6M). As mitochondrial mass was reduced by ZONAB depletion, the remaining mitochondria must thus work at a higher rate to achieve the same basal activity, which was also supported by the MitoTracker/TOMM20 ratios (Fig. 6K,M). Supporting mitochondrial function with nicotinamide, which attenuates senescence (Fig. 6D,E), rescued much of the ZONAB depletion-induced drop in compensatory respiration, and nicotinamide-treated control and ZONAB-depleted cells had the same maximal mitochondrial activity (Fig. 6L). Hence, mitochondrial dysfunction in ZONAB-depleted endothelial cells is a driver of the induction of cellular senescence.

**Figure 6.**
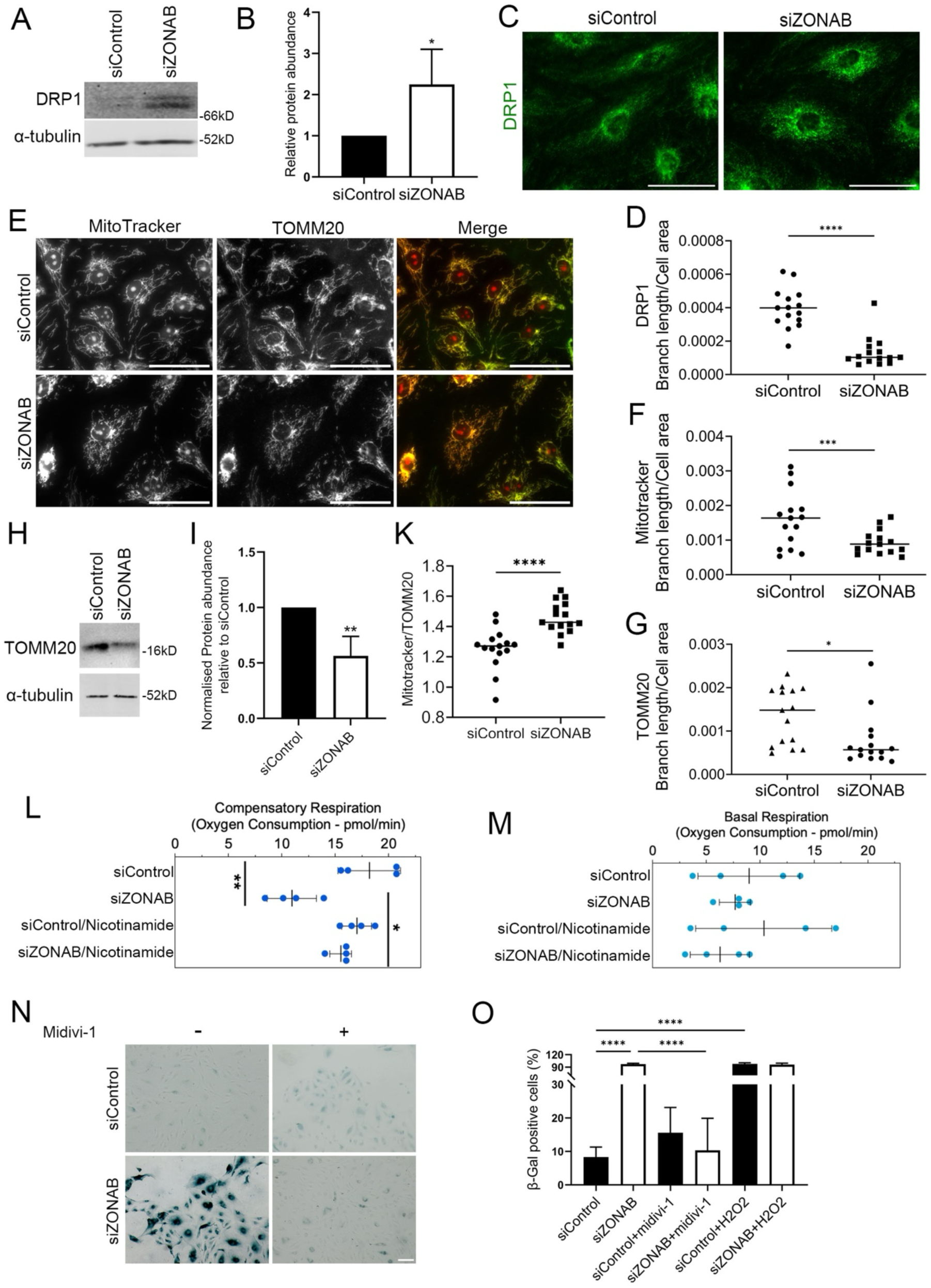
ZONAB deficiency leads to mitochondrial dysfunction. (A-B) Effect of ZONAB depletion on DRP1 expression assessed by immunoblotting. α-tubulin was used as a loading control in the densitometric quantification in panel B (means ± 1 SD, n=3). (C-D) Immunofluorescence staining of DRP1 indicates increased mitochondrial fragmentation (scale bar: 50μm). The quantification in panel D shows branch length relative to the cell area of the DRP1 staining (shown are means and datapoints, n=15). (E) Images of TOMM20 immunofluorescence and MitoTracker staining in control or ZONAB siRNA-transfected HDMECs (scale bar: 50 μm). Panels F and G show a quantification of the branch length for MitoTracker (F) and TOMM20 (G) relative to the cell area. Panel K shows the ratio of MitoTracker to TOMM20 staining. All three quantifications show means and data points, n=15 images from 3 independent experiments). (H-I) Immunoblotting of TOMM20. Panel I shows a densitometric quantification of the immunoblots using α-tubulin as a loading control for normalization. (L-M) Measurements of compensatory and basal respiration using a mitochondrial stress test protocol combined with a Seahorse analyzer (shown are means ± 1 SD and datapoints, n=4). (N-O) SAβG staining of control and ZOANB siRNA-transfected cells after a 48-hour incubation without or with DRP1 inhibitor midivi-1. Scale bar: 200 μm. β-gal positive cells were imaged by bright field microscopy and quantified (panel H, means ± 1 SD, n=3). Panels B,D,F,G,K: t-tests. Panels L,M,O: Tukey HSD. *p<0.05, **p<0.01, ***p<0.001, ****p<0.001

Increased DRP1 expression can lead to mitochondrial malfunction, increased ROS, and cellular senescence ^36,37^. DRP1 can be inhibited by the small molecule inhibitor midivi-1 ^38^. Hence, we tested whether DRP1 inhibition attenuated the entry into cell senescence of ZONAB-depleted cells. Figure 6N-O shows that midivi-1 strongly reduced the number of SAßG-positive cells. Thus, inhibition of DRP1 attenuates cellular senescence of ZONAB-depleted cells, further supporting the link between ZONAB depletion-induced deregulation of mitochondria and cellular senescence.

### ZONAB depletion induces DNA methylation of genes involved in cellular senescence

Changes in DNA methylation have been shown not only to establish but to maintain and stabilise the cellular senescent phenotype ^39,40^. DNA methylation is an epigenetic mechanism that contributes to silencing genes required for cell proliferation. To test if ZONAB depletion affects DNA methylation, we used bisulfite-converted genomic DNA from control and ZONAB-depleted endothelial cells to screen an Illumina genome methylation EPIC array. We detected hypermethylated cytosines in the PLK-1 gene promoter in ZONAB-depleted cells (Fig. 7A), in agreement with the reduced expression described in Figure 4. Additionally, we identified several other differentially DNA methylated regions in the promoters of genes required for proliferation, such as WRN Helicase Interacting Protein 1 (WRNIP1), methyltransferase-like protein 7A (METTL7A), and the cell cycle-regulating transcription factor forkhead box M1 (FOXM1) that were found to be downregulated in the gene expression analysis (Table S2, S9). Expression of genes such as FOXM1 is essential to prevent cellular senescence ^41^. Hypermethylation of the FOXM1, WRNIP1, and METTL7A promoters correlated with decreased expression at the protein and mRNA levels (Fig. 7B-D). Thus, ZONAB depletion led to increased DNA methylation of genes whose repression leads to cell cycle arrest and cellular senescence.

**Figure 7.**
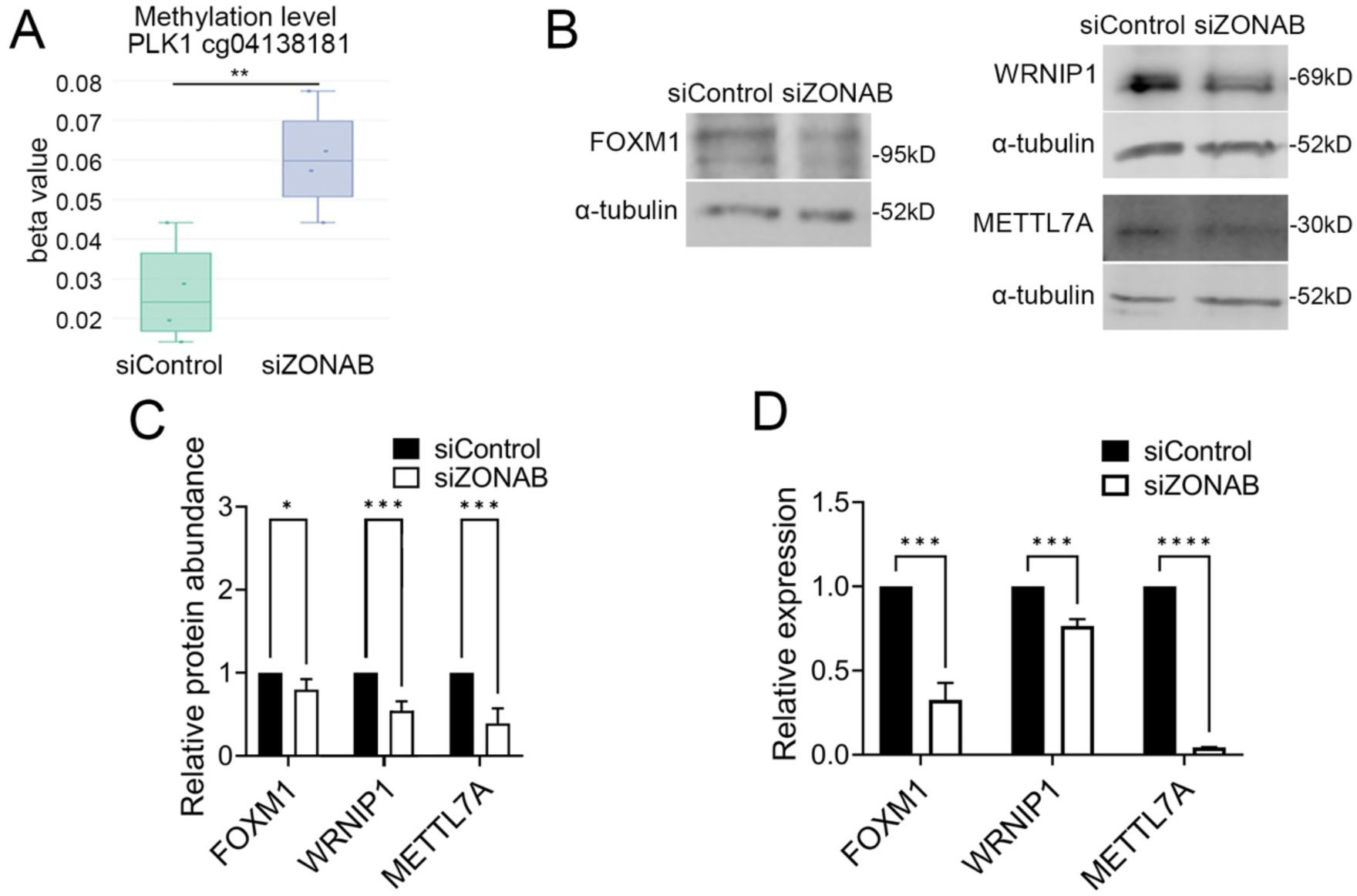
ZONAB regulates DNA methylation of genes involved in endothelial cell proliferation and senescence. (A) Boxplot analysis of the DNA methylation status of the PLK1 promoter at cg04138181 from the Illumina EPIC methylation array (n=4; median, center line; inter-quartile range, box; 95th percentiles, whisker). See Table S9 for examples of other genes identified as being differentially methylated. (B-C) Immunoblotting reveals repression of FOXM1, WRNIP1, and METTL7A. Densitometric quantification of at least 3 repeat experiments. Values were normalized to α-tubulin expression, and ratios for control siRNA-transfected samples were set to 1 (means ± 1 SD). (D) Analysis of mRNA expression of FOXM1, WRNIP1, and METTL7A relative to GAPDH determined by RT-qPCR (means ± 1 SD, n=3). T-tests: **p<0.01, ***p<0.001, ****p<0.001

## Discussion

Our data demonstrate that ZONAB regulates endothelial cell proliferation and migration, and that its expression is required to prevent endothelial cells from entering cellular senescence. Hence, ZONAB depletion was found to disrupt the normal angiogenic behaviour *in vitro* and growth factor-induced angiogenesis *in vivo*. On a cellular level, we found that ZONAB regulates the expression of cell cycle regulators that are required for key steps along the entire cell cycle, and that ZONAB depletion led to the induction of epigenetic changes on promoters of genes involved in cell proliferation and senescence. In ZONAB’s absence, senescence was induced by mitochondrial deregulation and a deficient oxidative stress response; hence, entry into the senescent state induced by ZONAB downregulation could be prevented by drugs impacting mitochondria, ROS levels, or PI3K activation.

ZONAB-depleted endothelial cells have a reduced angiogenic potential in culture and *in vivo*. The *in vitro* assay is quick and is primarily affected by the effect of ZONAB depletion on the cytoskeleton and cell migration; the *in vivo* assay spreads over days and is affected by defects in proliferation and migration. Senescent endothelial cells undergo a profound remodelling of the actin cytoskeleton, which includes increased F-actin stiffness, redistribution of focal adhesions, and elevated traction forces, leading to less mobile cells that form more stable focal adhesions ^42,43^. We observed similar cytoskeletal changes in ZONAB-depleted cells. Hence, the reduced migration is a likely reflection of increased stability of focal adhesions. As senescent cells also cease to proliferate, both the *in vitro* and *in vivo* angiogenesis defects are thus likely to be direct consequences of ZONAB-depleted cells having entered cellular senescence.

The cytoskeletal remodelling of ZONAB-depleted cells also affected junctional proteins, most notably JACOP, an actomyosin-organising adaptor, and claudin-5, a major tight junction adhesion protein required for blood-brain barrier integrity ^44^. Vinculin was removed from cell-cell junctions and redistributed to focal adhesions, which coincided with an analogous redistribution of active myosin-II and F-actin. Vinculin recruitment is a well-established indicator of increased tensile forces acting on focal adhesions and adherens junctions, respectively. Hence, the cytoskeletal phenotype indicates a reinforcement of cell-substrate adhesions and a weakening of adherens and tight junctions. Such changes have also been observed in senescent endothelial cells *in vivo*, as vascular cell senescence increases tight junction permeability and decreases blood-brain barrier function in aged mice ^45^.

Proliferation of ZONAB-depleted endothelial cells was strongly inhibited in agreement with the senescent phenotype. Cell cycle regulation was also observed in immortalised epithelial cell lines that did not become senescent; hence, ZONAB’s role in proliferation may lead to senescence but also regulates proliferation in non-senescent cells ^9,11,46^. We had previously identified cyclin D1 and PCNA as ZONAB-regulated genes in epithelial cells ^11^. Cyclin D1 was also strongly downregulated in endothelial cells, but the effect on PCNA was very modest. However, a range of major cell cycle genes were affected by ZONAB depletion, including cyclins and kinases affecting different steps of the cell cycle; hence, ZONAB depletion has a broad effect on cell cycle regulation.

Some of the new ZONAB-regulated cell cycle genes identified here are involved in the regulation of endothelial proliferation, angiogenesis and senescence. For example, cyclin A2 plays a crucial role in activating PLK-1, which is essential for cell proliferation ^47^. Both cyclin A2 and PLK-1 are downregulated during cell senescence ^48,49^, and silencing PLK-1 in endothelial cells reduces tube-formation, implicating PLK-1 in the regulation of angiogenic behaviour ^50^. Hence, both genes are ZONAB signalling targets important for angiogenesis.

It is unlikely that all the genes deregulated upon ZONAB depletion are direct targets of ZONAB-regulated transcription, as the cyclin D1 gene ^11^. We discovered here that ZONAB signalling regulates the methylation of some of the genes that are downregulated or upregulated upon its depletion. For instance, the *PLK-1* promoter, which is hypermethylated in ZONAB-depleted cells, also becomes hypermethylated in response to ROS, leading to reduced *PLK-1* expression and cell senescence ^51^. DNA methylation patterns in genes related to endothelial senescence are known to impact endothelial cell function and to contribute to vascular ageing and related diseases ^52^. Senescence-associated DNA methylation often involves global hypomethylation (decreased methylation across the genome) and focal hypermethylation (increased methylation at specific gene locations) ^53^. We identified several genes linked to cell proliferation and senescence that became hypermethylated in response to ZONAB depletion. This included genes associated with ageing, such as *WRNIP1*, *FOXM1*, and *METTL7A* ^54–56^. While *PARD3* induction was suggested to play a role as a biomarker for cell senescence ^57^. Our results thus uncover a novel role of ZONAB in regulating DNA methylation and, thereby, the regulation of genes important for entry into cell senescence.

While downregulation of the cell cycle is a normal feature of cell senescence, cell cycle inhibition is a process that generally does not lead to entry into senescence. Hence, ZONAB depletion in endothelial cells must induce changes not directly linked to the cell cycle. A common stimulus leading to cell senescence is increased ROS and oxidative stress. ZONAB depletion not only led to an increase in ROS but also to dysregulation of mitochondria, which is known to contribute to ROS induction. Our data further indicate that ZONAB regulates the expression of PGC-1α. PGC-1α regulates mitochondrial biosynthesis and, together with Nrf2, expression of HO-1. Hence, even in the presence of increased Nrf2 expression, ZONAB depletion led to HO-1 repression and, therefore, failed ROS elimination, triggering cellular senescence. ROS stimulates cell senescence via a p53/p21-dependent mechanism ^3,24,25^; ZONAB depletion indeed stimulated expression of both p53 and p21.

ROS induces PI3K/Akt ^26^. We have observed a strong increase in phosphorylation of the PI3K substrate site of Akt, and an inhibition of senescence in PI3K inhibitor-treated ZONAB-depleted endothelial cells. The most studied inducer of cell senescence is H2O2, which directly increases ROS and which can be rescued by Nicotinamide or NAC ^58,59^. With both drugs, we were also able to rescue ZONAB-depletion-induced senescence in endothelial cells, further supporting the conclusion that the increase in ROS was responsible for the induction of senescence of ZONAB-depleted endothelial cells.

In addition to the repression of HO-1 and PGC-1α, we observed a reduction in mitochondrial mass and total respiratory capacity, which required remaining mitochondria to be more active. Mitochondrial dysregulation stimulates ROS production. ZONAB depletion led to increased expression of DRP-1 and, consequently, increased mitochondrial fragmentation. Entry into senescence was inhibited by a DRP-1 inhibitor or by stimulating mitochondrial function with nicotinamide. Hence, the regulation of mitochondria downstream of ZONAB signalling plays a functionally important role in endothelial cell homeostasis.

Endothelial cell senescence is thought to contribute to the development and/or progression of diseases such as atherosclerosis and microvascular conditions affecting the heart or brain, as well as neurodegenerative disorders such as Alzheimer’s disease ^1,2,60^. Genome-wide genetic studies have shown that the ZONAB gene is associated with loci with single-nucleotide polymorphisms that increase the risk for developing atherosclerosis as well as high systolic blood pressure ^4–6^. While it is currently unknown if and how those SNPs affect ZONAB function and/or expression, these observations, combined with our data, suggest that ZONAB malfunction may drive serious vascular diseases by inducing cellular senescence. Similarly, a recent study identified ZONAB/YBX3 as a transcription factor that interacts with enhancer sequence elements of alleles identified as risk loci for Alzheimer’s disease ^61^. Hence, disruption of the here-identified role of ZONAB in regulating endothelial angiogenic behaviour and cellular senescence may promote common vascular and neurodegenerative diseases. Establishment of the necessary disease models will be required to test the importance of ZONAB or, rather, its downregulation in the development of such endothelial diseases.

## Resource Availability

### Lead Contact

Requests for further information and resources should be directed to and will be fulfilled by the lead contact, Maria S. Balda

### Materials Availability

This study did not generate new unique reagents.

### Data and Code Availability

The RNA sequencing and expression array data will be made available on the European Nucleotide Archive. All other data are available from the contact.

## Acknowledgments

We thank Dr Olga Tornavaca for the studies of ZONAB and angiogenesis; unfortunately, we were unable to contact her to read the current manuscript. We are grateful to the IoO Imaging Unit for their support and assistance during this work.

This work was supported by the British Heart Foundation (CRM:0001059), the Macular Disease Society (22-RG-3), the Moorfields Eye Charity (R180018A), the MRC (MR/W028735/1 and G0900098), BBSRC (BB/X000575/1), and the NIHR Biomedical Research Centre at Moorfields Eye Hospital.

## Author Contributions

WJ, EL, and JD performed the majority of the reported experiments. The angiogenesis experiments were done by Olga Tornavaca, GMB, and AMR. EL performed the methylation experiments. KM and MSB carried out the expression analysis by cDNA array and RNA sequencing. All authors contributed to data curation and formal analysis. KM and MSB conceived the study and methodology, were responsible for funding acquisition, project administration and supervision, performed investigation, data curation, and formal analysis, and wrote the manuscript with the support of all authors.

## MATERIALS AND METHOD

### Cell culture and transfection

Human dermal microvascular endothelial cells (HDMEC adult C-12212 and juvenile C-12210) were obtained from Promo Cell, plated on 0.5% gelatine-coated tissue culture dishes in Medium MV2 with C-39225 supplement mix (ECGMv2, Promo Cell). Cells were used between passages two and six. C57BL/6 mice were used for the Matrigel plug assays. HDMECs were transfected with a pool or single siRNAs against ZONAB (5’-AGACGUGGCUACUAUGGAA-3’ and 5’-CAACGUCAGAAAUGGAUAU-3’) or nontargeting siRNAs (5’-UGGUUUACAUGUCGACUAA and 5’-UGGUUUACAUGUUGUGUGA-3’). All siRNAs were obtained from Sigma-Aldrich and were synthesised with dTdT 3’-overhangs. Depending on the experimental condition, HDMECs were plated in different types of dishes (e.g., 30,000 cells on 10-mm cover glasses in 48-well dishes, cover glass coated with 0.8 mg/ml Growth Factor Reduced Matrigel, Corning 354230; or 6-well dishes, 200000 cells per well). The following day, transfection of siRNAs was performed using Lipofectamine RNAmax according to the manufacturer’s instructions. The medium was replaced 24 hours later, and the cells were collected after 2–3 days of additional culture for further analysis.

### Immunofluorescence and protein analysis

Cells were fixed either with methanol at −20°C for 10 min or with 3% PFA for 20 min at room temperature. PFA-fixed cells were permeabilised with 0.3% Triton X-100 in PBS containing 0.3% BSA for 5 min. The cells were incubated overnight at 4°C with primary antibodies. All primary antibodies used are listed in Table S10. The samples were then washed three times with PBS before incubating with fluorescent secondary antibodies (Table S10). For nuclei visualisation, Hoechst 33258 (Sigma-Aldrich) was used. Cells on coverslips were mounted with Prolong Gold mounting medium (Thermofisher Scientific) and stored at 4°C. Images were taken with a fluorescent microscope (Zeiss Axio Skop MOT2 microscope) with a 40x/NA1.2 or 60x/NA1.4 oil immersion objectives using Simple PCI software (Hamamatsu Photonics) or an inverted Nikon microscope (Eclipse Ti-E) using a 60x/1.4 NA oil immersion objective lens, a CoolSNAP HQ2 camera (Photometrics), and Nikon software. Adobe Photoshop or Fuji/ImageJ was used to adjust contrast. For immunoblotting of whole-cell lysates, the cells were washed twice with PBS, lysed in SDS-PAGE sample buffer, and denatured at 70°C for 10 min ^11^. The samples were then homogenised with a 1 ml syringe and a 25- and 34g needle before running on SDS-PAGE gels and transferring to PVDF or nitrocellulose membranes. The membranes were washed and blocked with 5% defatted milk powder dissolved in PBS containing 0.1% Tween-20. The used primary and secondary antibodies are detailed in Table S10.

### Matrigel plug angiogenesis assay

The Matrigel plug assay was performed and analysed as described previously (Birdsey et al., 2008). In brief, C57BL/6 mice were injected with a mix of Matrigel (BD), heparin, and, depending on the condition, FGF, and 2 μM siRNAs near the abdominal midline. Plugs were harvested after 7 days postmortem, fixed in 4% paraformaldehyde in PBS for 2 hours at room temperature, transferred to 70% ethanol, embedded in paraffin, and processed for haematoxylin and eosin staining. The same siRNAs were used as for human cells (the corresponding sequences between human and mouse are 100% identical). For quantification, vessels contained in the Matrigel plug were identified by the presence of nucleated cells surrounding a lumen containing red blood cells. Vessels were counted in four fields of view using a 20× objective lens. Experiments were performed according to the Animals (Scientific Procedures) Act of 1986.

### Capillary-like formation on Matrigel and cell migration

Ice-cold growth factor-reduced Matrigel (BD Biosciences; 8–10 mg/ml, 50 μl/well) was added to flat-bottom 96-well plates and allowed to solidify for 1 hour at 37°C. HDMECs (15,000 cells/well) that had been transfected with siRNAs 48 hours before were seeded in duplicates into the coated wells. The plates were then incubated at 37°C in ECGMv2. Pictures were taken at different times, and capillaries were fixed in phosphate-buffered formalin at the end of the experiment. Branching points were quantified in four different fields and two different experiments per condition. For cell migration analysis, HDMECs were plated in 48-well dishes, and siRNAs were transfected as described above. 40 hours later, a scratch wound was inflicted with a plastic yellow tip, and pictures were taken (zero timepoint). Cells were then allowed to migrate at 37°C. Pictures were taken at different times using a 5×/0.12 NA objective lens on an inverted microscope (DMIRB; Leica), a camera (C4742-95; Hamamatsu Photonics), and simple PCI software (ambient temperature). The area without cells was calculated at the different time points using ImageJ.

### Bromodeoxyuridine incorporation assay

HDMECs were cultured on glass coverslips and transfected with siRNAs. The ells were starved for 2 days to synchronise proliferation by using medium with one-tenth of the supplement. Complete ECGMv2 medium containing Bromodeoxyuridine (BrdU, 10 μM) was then added, and the cells were incubated for 4 or 8 hours. The cells were then fixed in 100% methanol for 7 minutes at -20°C, washed twice with PBS, and incubated with 1.5M HCL for 30 minutes. After washing 4 times with PBS and blocking with 1% BSA and 0.1% NaN3 in PBS overnight, the cells were incubated with mouse antibodies specific for BrdU and DNase1 for 1 hour. After washing three times with 1% BSA and 0.1 % NaN3 in PBS, the cells were incubated with fluorescent secondary antibody and Hoechst dye for one hour. The samples were then washed and mounted onto glass slides with ProLong® Gold Antifade Reagent. BrdU-positive cells were counted and quantified as a percentage of all cells per field of view.

### Cell Trace CSFE

The day after RNA interference, the supplement of the medium was reduced to one-tenth, and, 24 hours later, Cell Trace-DMSO stock was added at a final concentration of 1 μM for 20 minutes. Cells were then washed with complete ECGMv2 medium. Control cells for the baseline were collected immediately, and samples for measuring the effect on proliferation were collected 48 hours later. Cells were collected in Trypsin EDTA, spun down at 1500rpm for 5 minutes, and fixed in 3% PFA for 15 minutes. CSFE levels were measured by flow cytometry using a BD LSRFortressa TM machine.

### Cell number

To assess cell number, DNA content was measured. In brief, cells were plated in triplicate into 96-well plates at equal cell densities, and RNA inference was performed as described above. The medium was removed after three days, and samples were frozen at −80°C. DNA contents were quantified using a CyQUANT Cell Proliferation Assay kit (Thermo Fisher Scientific) according to the manufacturer’s instructions.

### Propidium iodide assay

Cell cycle analysis was performed by flow cytometry using propidium iodide. Cells were collected 72h after transfection using trypsin for 10 minutes. They were then centrifuged and rinsed twice in PBS before fixation with cold 70% ethanol for 30 minutes on ice. Following fixation, the cells were centrifuged and rinsed again twice with PBS. The PBS was removed after the final rinse, and 50 μl of RNase A solution in PBS (Roche 10109142001, 100 μg/ml, DNAse-free) was added to the pellet, followed by 400 μl of propidium iodide solution in PBS (Sigma P4170, 50 μg/ml). After a 5-minute incubation at room temperature, the propidium iodide staining was measured using a BD LSRFortessa™ X-20 Cell Analyzer with BD FACS Diva software (BD Biosciences). Data analysis was performed using Flowing Software (Perttu Terho, Turku Centre for Biotechnology). At least 10,000 events (single cells positive for PI staining) were analysed per sample.

### Quantitative polymerase chain reaction (qPCR)

On ice, cDNA (50 ng/μl) was placed in quadruplicates into a MicroAmp Optical 96-well Reaction Plate (Thermo Fisher Scientific) with PowerUp SYBR Green Master Mix (Thermo Fisher Scientific) and 5 μM each of the two primers of interest (detailed in Table S11). A reaction volume of 10 μl was used. Fluorescence was measured using a Quant Studio 6 Flex Real-Time PCR System (Thermo Fisher Scientific). Relative mRNA levels were determined by standardising target primer fluorescence signals with the corresponding endogenous control fluorescence signals. Values were then normalised with the respective control values (Quant Studio 6 software was used).

### Senescence-associated β-galactosidase assay

30,000 cells per well were plated in 48-well plates coated with 0.8 ml/ml Matrigel GFR. The following day, siRNA transfections were performed. After 3 days, the cells were washed in PBS, fixed in 3% paraformaldehyde (5 minutes, room temperature), washed again in PBS and incubated at 37°C overnight (without CO₂) with senescence-associated β-Galactosidase staining solution (0.1% 5-bromo-4-chloro-3-indolyl b-D-galactosidase (X-Gal), 5mM potassium ferrocyanide, 5 mM potassium ferricyanide, 150 mM NaCl, 2 mM MgCl_2_ in 40 mM citric acid/sodium phosphate, pH 6.0). For positive (lysosomal β-gal) and negative (bacterial β-gal) controls, the solution was brought to pH 4.0 or pH 7.5, respectively. After colour development, the cells were washed twice with PBS and once with methanol before examination with a brightfield Leica DM IL microscope fitted with a 20x objective. At least three images were captured at different locations for each coverslip, and senescence was determined by the percentage of SA-β-gal-positive cells per image.

### Caspase and necrosis assays

The activities of caspases 3 and 7 were determined using the Apo-One® Homogeneous Caspase 3/7 assay kit (Promega, Madison, WI) according to the manufacturer’s instructions. The fluorescence of the samples was measured at 490 nm excitation and 530 nm emission with a FLUOstar Optima plate reader (BMG LABTECH Inc., Durham, NC). Necrosis was assessed by measuring lactate dehydrogenase release into the medium using the CytoTox-ONE^TM^ Homogeneous Membrane Integrity Assay (Promega, Madison, WI), according to the manufacturer’s recommendations. The fluorescence of the samples was measured at 560 nm excitation and 590 nm emission with the FLUOstar Optima plate reader.

### ROS Assays

Two sets of cells were plated in a 96-well plate in quadruplicate samples for each condition. The cells were then transfected with control or ZONAB siRNAs. After 3 days, one set of samples received MitoTracker Red CMXRos, which becomes fluorescent when oxidised. The second set of samples received MitoTracker Green FM. Both MitoTrackers were added at a concentration of 100 nM and were incubated for 10 minutes. Cells were washed in medium and then left for 10 minutes in fresh medium. Cells were washed again in medium to remove unincorporated MitoTracker reagents before fixation with 3% PFA in PBS. Fluorescence was then measured using a FLUOstar OPTIMA plate reader and BMG LABTECH software (MitoTracker Red CMXRos, excitation 540 nm and emission 590 nm; MitoTracker Green FM, excitation 485 nm, emission 525 nm).

### Seahorse Assays

Cells were seeded into a Seahorse XFe96/XF Pro Cell Culture Microplate (Agilent, USA) at a density of 0.7*10^4^ cells/well in 50 μl of ECGMv2 medium. The following day, the cells were transfected with siRNAs for 24 hours followed by incubating in the fresh culture medium for 48-72 hours. One hour before the Seahorse assay, the culture medium was replaced with Seahorse XF Assay Medium, composed of Seahorse XF DMEM Medium (pH 7.4, Agilent, USA) supplemented with 10 mM glucose, 1 mM pyruvate, 2 mM L-glutamine, and 1% C-39225 supplement (Agilent, USA). All experiments measuring energy metabolism were run with a Seahorse XF Pro Analyzer using the software to run and analyse the experiments provided by the manufacturer (Agilent, USA). The mitochondrial stress test was run according to the manufacturer’s instructions using reagents purchased from Merck Life Science UK. Assay results were normalised by quantifying cell numbers using the CyQUANT assay. Briefly, cells in the Seahorse Cell Culture Microplates were frozen overnight at -80°C. The CyQUANT assay was performed using the CyQUANT™ Cell Proliferation Assay Kit (Invitrogen, USA) the next day, following the manufacturer’s guidelines. Fluorescence was measured at an excitation wavelength of 480 nm and an emission wavelength of 520 nm. The CyQUANT data were uploaded into the Seahorse Wave software for normalisation of the Seahorse experiments.

### Expression analysis by Affymetrix cDNA gene expression microarray and RNA sequencing

Cells plated in 6-well dishes were transfected with either control or ZONAB siRNAs. After 4 days, total RNA was isolated using the RNeasy Mini Kit (Qiagen). The RNA samples were then analysed by UCL Genomics using either an Affymetrix cDNA gene expression microarray or by RNA sequencing. Labelled gene chips were scanned, using a confocal argon ion laser (Agilent Technologies), and the data were analysed using Gene Spring 7.2 software (Agilent). RNA sequencing was also performed by UCL Genomics. Briefly, libraries were generated using a Watchmaker RNA sequencing kit with mRNA enrichment and unique dual indexing. Sequencing was performed with an Illumina NextSeq 2000 system. The data was analysed using the Galaxy Europe site (https://usegalaxy.eu/) for the differential expression analysis (with MultiQC, RNA STAR, htseq-count and DESeq2) using the Ensembl GRCh37 (hg19) annotation file and Galaxy ProteoRE (https://proteore.org/) for the functional and pathway analysis ^62–69^.

### DNA methylation arrays

Cells plated in 6-well dishes were transfected with siRNAs the following day. After 4 days, genomic DNA from cells was extracted using a QIA amp DNA Mini and Blood Mini kit (Qiagen) according to manufacturer instructions. To identify methylated or unmethylated cytosines, DNA was treated with sodium bisulfite using the EZ DNA methylation kit (Zymo Research, CA, USA) according to kit instructions. An Infinium® Methylation EPIC Bead Chip was used for the analysis at UCL Genomics Facility, using 45 μl of each sample containing 500ng of DNA according to the kit instructions. Analysis was completed either with Illumina Genome Studio software or with the R programme Bioconductor package, the Chip Analysis Methylation Pipeline (Champ). Using this package’s samples.idat, the data was loaded, normalised, and analysed for singular value decomposition (SVD) and statistically significant methylated regions (DMRs) were found as described (Morris et al. 2014).

### Image Analysis and Statistics

All image analysis and quantification of immunoblots were performed with either Image J/Fiji or Adobe Photoshop. Quantitative data are shown as datapoints and/or means +/-one standard deviation. The n number is provided in the respective figure legends and refers to mice in the *in vivo* studies in Figure 1 or independent biological repeat experiments in the cell-based assays, except otherwise indicated in the figure legends. Statistical analysis was performed with JMP Pro or GraphPad Prism. Statistical significance of data pairs was assessed with two-sided t-tests and, in experiments requiring multiple comparisons, Tukey-Kramer HSD tests following an ANOVA test.

## References

1. Georgieva, I., Tchekalarova, J., Iliev, D., and Tzoneva, R. (2023). Endothelial Senescence and Its Impact on Angiogenesis in Alzheimer’s Disease. Int J Mol Sci 24. 10.3390/ijms241411344.

2. Han, Y., and Kim, S.Y. (2023). Endothelial senescence in vascular diseases: current understanding and future opportunities in senotherapeutics. Exp Mol Med 55, 1–12. 10.1038/s12276-022-00906-w.

3. Huang, W., Hickson, L.J., Eirin, A., Kirkland, J.L., and Lerman, L.O. (2022). Cellular senescence: the good, the bad and the unknown. Nat Rev Nephrol 18, 611–627. 10.1038/s41581-022-00601-z.

4. Koyama, S., Ito, K., Terao, C., Akiyama, M., Horikoshi, M., Momozawa, Y., Matsunaga, H., Ieki, H., Ozaki, K., Onouchi, Y., et al. (2020). Population-specific and trans-ancestry genome-wide analyses identify distinct and shared genetic risk loci for coronary artery disease. Nat Genet 52, 1169–1177. 10.1038/s41588-020-0705-3.

5. Aragam, K.G., Jiang, T., Goel, A., Kanoni, S., Wolford, B.N., Atri, D.S., Weeks, E.M., Wang, M., Hindy, G., Zhou, W., et al. (2022). Discovery and systematic characterization of risk variants and genes for coronary artery disease in over a million participants. Nat Genet 54, 1803–1815. 10.1038/s41588-022-01233-6.

6. Verma, A., Huffman, J.E., Rodriguez, A., Conery, M., Liu, M., Ho, Y.L., Kim, Y., Heise, D.A., Guare, L., Panickan, V.A., et al. (2024). Diversity and scale: Genetic architecture of 2068 traits in the VA Million Veteran Program. Science 385, eadj1182. 10.1126/science.adj1182.

7. Jiang, L., Zheng, Z., Fang, H., and Yang, J. (2021). A generalized linear mixed model association tool for biobank-scale data. Nat Genet 53, 1616–1621. 10.1038/s41588-021-00954-4.

8. Balda, M.S., and Matter, K. (2000). The tight junction protein ZO-1 and an interacting transcription factor regulate ErbB-2 expression. Embo j 19, 2024–2033. 10.1093/emboj/19.9.2024.

9. Balda, M.S., Garrett, M.D., and Matter, K. (2003). The ZO-1-associated Y-box factor ZONAB regulates epithelial cell proliferation and cell density. J Cell Biol 160, 423–432. 10.1083/jcb.200210020.

10. Lima, W.R., Parreira, K.S., Devuyst, O., Caplanusi, A., N’Kuli, F., Marien, B., Van Der Smissen, P., Alves, P.M., Verroust, P., Christensen, E.I., et al. (2010). ZONAB promotes proliferation and represses differentiation of proximal tubule epithelial cells. J Am Soc Nephrol 21, 478–488. 10.1681/asn.2009070698.

11. Sourisseau, T., Georgiadis, A., Tsapara, A., Ali, R.R., Pestell, R., Matter, K., and Balda, M.S. (2006). Regulation of PCNA and cyclin D1 expression and epithelial morphogenesis by the ZO-1-regulated transcription factor ZONAB/DbpA. Mol Cell Biol 26, 2387–2398. 10.1128/MCB.26.6.2387-2398.2006.

12. Tsapara, A., Matter, K., and Balda, M.S. (2006). The heat-shock protein Apg-2 binds to the tight junction protein ZO-1 and regulates transcriptional activity of ZONAB. Mol Biol Cell 17, 1322–1330. 10.1091/mbc.e05-06-0507.

13. Nie, M., Balda, M.S., and Matter, K. (2012). Stress- and Rho-activated ZO-1-associated nucleic acid binding protein binding to p21 mRNA mediates stabilization, translation, and cell survival. Proc Natl Acad Sci U S A 109, 10897–10902. 10.1073/pnas.1118822109.

14. Frankel, P., Aronheim, A., Kavanagh, E., Balda, M.S., Matter, K., Bunney, T.D., and Marshall, C.J. (2005). RalA interacts with ZONAB in a cell density-dependent manner and regulates its transcriptional activity. EMBO J 24, 54–62. 10.1038/sj.emboj.7600497.

15. Dupasquier, S., Delmarcelle, A.S., Marbaix, E., Cosyns, J.P., Courtoy, P.J., and Pierreux, C.E. (2014). Validation of housekeeping gene and impact on normalized gene expression in clear cell renal cell carcinoma: critical reassessment of YBX3/ZONAB/CSDA expression. BMC Mol Biol 15, 9. 10.1186/1471-2199-15-9.

16. Pannequin, J., Delaunay, N., Darido, C., Maurice, T., Crespy, P., Frohman, M.A., Balda, M.S., Matter, K., Joubert, D., Bourgaux, J.F., et al. (2007). Phosphatidylethanol accumulation promotes intestinal hyperplasia by inducing ZONAB-mediated cell density increase in response to chronic ethanol exposure. Mol Cancer Res 5, 1147–1157. 10.1158/1541-7786.MCR-07-0198.

17. Qin, H.L., Wang, X.J., Yang, B.X., Du, B., and Yun, X.L. (2021). Notoginsenoside R1 attenuates breast cancer progression by targeting CCND2 and YBX3. Chin Med J (Engl) 134, 546–554. 10.1097/CM9.0000000000001328.

18. Wang, C., You, Z., He, Y., and Chen, X. (2023). Identification of RNA-binding protein YBX3 as an oncogene in clear cell renal cell carcinoma. Funct Integr Genomics 23, 225. 10.1007/s10142-023-01145-6.

19. Sun, Y., Li, Z., Wang, W., Zhang, X., Li, W., Du, G., Yin, J., Xiao, W., and Yang, H. (2022). Identification and verification of YBX3 and its regulatory gene HEIH as an oncogenic system: A multidimensional analysis in colon cancer. Front Immunol 13, 957865. 10.3389/fimmu.2022.957865.

20. Tornavaca, O., Chia, M., Dufton, N., Almagro, L.O., Conway, D.E., Randi, A.M., Schwartz, M.A., Matter, K., and Balda, M.S. (2015). ZO-1 controls endothelial adherens junctions, cell-cell tension, angiogenesis, and barrier formation. J Cell Biol 208, 821–838. 10.1083/jcb.201404140.

21. Normandin, K., Coulombe-Huntington, J., St-Denis, C., Bernard, A., Bourouh, M., Bertomeu, T., Tyers, M., and Archambault, V. (2023). Genetic enhancers of partial PLK1 inhibition reveal hypersensitivity to kinetochore perturbations. PLoS Genet 19, e1010903. 10.1371/journal.pgen.1010903.

22. Astle, M.V., Hannan, K.M., Ng, P.Y., Lee, R.S., George, A.J., Hsu, A.K., Haupt, Y., Hannan, R.D., and Pearson, R.B. (2012). AKT induces senescence in human cells via mTORC1 and p53 in the absence of DNA damage: implications for targeting mTOR during malignancy. Oncogene 31, 1949–1962. 10.1038/onc.2011.394.

23. Chan, K.T., Blake, S., Zhu, H., Kang, J., Trigos, A.S., Madhamshettiwar, P.B., Diesch, J., Paavolainen, L., Horvath, P., Hannan, R.D., et al. (2020). A functional genetic screen defines the AKT-induced senescence signaling network. Cell Death Differ 27, 725–741. 10.1038/s41418-019-0384-8.

24. Mijit, M., Caracciolo, V., Melillo, A., Amicarelli, F., and Giordano, A. (2020). Role of p53 in the Regulation of Cellular Senescence. Biomolecules 10. 10.3390/biom10030420.

25. Terao, R., Ahmed, T., Suzumura, A., and Terasaki, H. (2022). Oxidative Stress-Induced Cellular Senescence in Aging Retina and Age-Related Macular Degeneration. Antioxidants (Basel) 11. 10.3390/antiox11112189.

26. Koundouros, N., and Poulogiannis, G. (2018). Phosphoinositide 3-Kinase/Akt Signaling and Redox Metabolism in Cancer. Front Oncol 8, 160. 10.3389/fonc.2018.00160.

27. Kasai, S., Shimizu, S., Tatara, Y., Mimura, J., and Itoh, K. (2020). Regulation of Nrf2 by Mitochondrial Reactive Oxygen Species in Physiology and Pathology. Biomolecules 10. 10.3390/biom10020320.

28. Luo, W., Li, J., Li, Z., Lin, T., Zhang, L., Yang, W., Mai, Y., Liu, R., Chen, M., Dai, C., et al. (2021). HO-1 nuclear accumulation and interaction with NPM1 protect against stress-induced endothelial senescence independent of its enzymatic activity. Cell Death Dis 12, 738. 10.1038/s41419-021-04035-6.

29. Luo, W., Wang, Y., Yang, H., Dai, C., Hong, H., Li, J., Liu, Z., Guo, Z., Chen, X., He, P., et al. (2018). Heme oxygenase-1 ameliorates oxidative stress-induced endothelial senescence via regulating endothelial nitric oxide synthase activation and coupling. Aging (Albany NY) 10, 1722–1744. 10.18632/aging.101506.

30. D’Egidio, F., Qosja, E., Ammannito, F., Topi, S., d’Angelo, M., Cimini, A., and Castelli, V. (2025). Antioxidant and Anti-Inflammatory Defenses in Huntington’s Disease: Roles of NRF2 and PGC-1alpha, and Therapeutic Strategies. Life (Basel) 15. 10.3390/life15040577.

31. Jain, A., Lamark, T., Sjottem, E., Larsen, K.B., Awuh, J.A., Overvatn, A., McMahon, M., Hayes, J.D., and Johansen, T. (2010). p62/SQSTM1 is a target gene for transcription factor NRF2 and creates a positive feedback loop by inducing antioxidant response element-driven gene transcription. J Biol Chem 285, 22576–22591. 10.1074/jbc.M110.118976.

32. Fernandez-Marcos, P.J., and Auwerx, J. (2011). Regulation of PGC-1alpha, a nodal regulator of mitochondrial biogenesis. Am J Clin Nutr 93, 884S–890. 10.3945/ajcn.110.001917.

33. Uchikado, Y., Ikeda, Y., and Ohishi, M. (2022). Current Understanding of the Pivotal Role of Mitochondrial Dynamics in Cardiovascular Diseases and Senescence. Front Cardiovasc Med 9, 905072. 10.3389/fcvm.2022.905072.

34. Miwa, S., Kashyap, S., Chini, E., and von Zglinicki, T. (2022). Mitochondrial dysfunction in cell senescence and aging. J Clin Invest 132. 10.1172/jci158447.

35. Kim, Y.Y., Um, J.H., Yoon, J.H., Lee, D.Y., Lee, Y.J., Kim, D.H., Park, J.I., and Yun, J. (2020). p53 regulates mitochondrial dynamics by inhibiting Drp1 translocation into mitochondria during cellular senescence. Faseb j 34, 2451–2464. 10.1096/fj.201901747RR.

36. Araujo, A.P.B., Vargas, G., Hayashide, L.S., Matias, I., Andrade, C.B.V., de Carvalho, J.J., Gomes, F.C.A., and Diniz, L.P. (2024). Aging promotes an increase in mitochondrial fragmentation in astrocytes. Front Cell Neurosci 18, 1496163. 10.3389/fncel.2024.1496163.

37. Kaur, S., Khullar, N., Navik, U., Bali, A., Bhatti, G.K., and Bhatti, J.S. (2024). Multifaceted role of dynamin-related protein 1 in cardiovascular disease: From mitochondrial fission to therapeutic interventions. Mitochondrion 78, 101904. 10.1016/j.mito.2024.101904.

38. Manczak, M., Kandimalla, R., Yin, X., and Reddy, P.H. (2019). Mitochondrial division inhibitor 1 reduces dynamin-related protein 1 and mitochondrial fission activity. Hum Mol Genet 28, 177–199. 10.1093/hmg/ddy335.

39. Cruickshanks, H.A., McBryan, T., Nelson, D.M., Vanderkraats, N.D., Shah, P.P., van Tuyn, J., Singh Rai, T., Brock, C., Donahue, G., Dunican, D.S., et al. (2013). Senescent cells harbour features of the cancer epigenome. Nat Cell Biol 15, 1495–1506. 10.1038/ncb2879.

40. Crouch, J., Shvedova, M., Thanapaul, R., Botchkarev, V., and Roh, D. (2022). Epigenetic Regulation of Cellular Senescence. Cells 11. 10.3390/cells11040672.

41. Smirnov, A., Panatta, E., Lena, A., Castiglia, D., Di Daniele, N., Melino, G., and Candi, E. (2016). FOXM1 regulates proliferation, senescence and oxidative stress in keratinocytes and cancer cells. Aging (Albany NY) 8, 1384–1397. 10.18632/aging.100988.

42. Cheung, T.M., Yan, J.B., Fu, J.J., Huang, J., Yuan, F., and Truskey, G.A. (2015). Endothelial Cell Senescence Increases Traction Forces due to Age-Associated Changes in the Glycocalyx and SIRT1. Cell Mol Bioeng 8, 63–75. 10.1007/s12195-014-0371-6.

43. Ferrari, S., and Pesce, M. (2021). Stiffness and Aging in Cardiovascular Diseases: The Dangerous Relationship between Force and Senescence. Int J Mol Sci 22. 10.3390/ijms22073404.

44. Nitta, T., Hata, M., Gotoh, S., Seo, Y., Sasaki, H., Hashimoto, N., Furuse, M., and Tsukita, S. (2003). Size-selective loosening of the blood-brain barrier in claudin-5-deficient mice. J Cell Biol 161, 653–660. 10.1083/jcb.200302070.

45. Yamazaki, Y., Baker, D.J., Tachibana, M., Liu, C.C., van Deursen, J.M., Brott, T.G., Bu, G., and Kanekiyo, T. (2016). Vascular Cell Senescence Contributes to Blood-Brain Barrier Breakdown. Stroke 47, 1068–1077. 10.1161/strokeaha.115.010835.

46. Zihni, C., Mills, C., Matter, K., and Balda, M.S. (2016). Tight junctions: from simple barriers to multifunctional molecular gates. Nat Rev Mol Cell Biol 17, 564–580. 10.1038/nrm.2016.80.

47. Silva Cascales, H., Burdova, K., Middleton, A., Kuzin, V., Müllers, E., Stoy, H., Baranello, L., Macurek, L., and Lindqvist, A. (2021). Cyclin A2 localises in the cytoplasm at the S/G2 transition to activate PLK1. Life Sci Alliance 4. 10.26508/lsa.202000980.

48. Kandhaya-Pillai, R., Miro-Mur, F., Alijotas-Reig, J., Tchkonia, T., Schwartz, S., Kirkland, J.L., and Oshima, J. (2023). Key elements of cellular senescence involve transcriptional repression of mitotic and DNA repair genes through the p53-p16/RB-E2F-DREAM complex. Aging (Albany NY) 15, 4012–4034. 10.18632/aging.204743.

49. Kim, H.J., Cho, J.H., and Kim, J.R. (2013). Downregulation of Polo-like kinase 1 induces cellular senescence in human primary cells through a p53-dependent pathway. J Gerontol A Biol Sci Med Sci 68, 1145–1156. 10.1093/gerona/glt017.

50. Gomes, C.P., Gomes-da-Silva, L.C., Ramalho, J.S., de Lima, M.C., Simões, S., and Moreira, J.N. (2013). Impact of PLK-1 silencing on endothelial cells and cancer cells of diverse histological origin. Curr Gene Ther 13, 189–201. 10.2174/1566523211313030004.

51. Ward, A., and Hudson, J.W. (2014). p53-Dependent and cell specific epigenetic regulation of the polo-like kinases under oxidative stress. PLoS One 9, e87918. 10.1371/journal.pone.0087918.

52. Xu, H., Li, S., and Liu, Y.S. (2021). Roles and Mechanisms of DNA Methylation in Vascular Aging and Related Diseases. Front Cell Dev Biol 9, 699374. 10.3389/fcell.2021.699374.

53. Sun, X., and Feinberg, M.W. (2021). Vascular Endothelial Senescence: Pathobiological Insights, Emerging Long Noncoding RNA Targets, Challenges and Therapeutic Opportunities. Front Physiol 12, 693067. 10.3389/fphys.2021.693067.

54. Yoshimura, A., Seki, M., and Enomoto, T. (2017). The role of WRNIP1 in genome maintenance. Cell Cycle 16, 515–521. 10.1080/15384101.2017.1282585.

55. Macedo, J.C., Vaz, S., Bakker, B., Ribeiro, R., Bakker, P.L., Escandell, J.M., Ferreira, M.G., Medema, R., Foijer, F., and Logarinho, E. (2018). FoxM1 repression during human aging leads to mitotic decline and aneuploidy-driven full senescence. Nat Commun 9, 2834. 10.1038/s41467-018-05258-6.

56. Shentu, T.P., Wu, T., Zhou, Z., Yeh, C.F., Zhu, J., Li, J., Miao, B.A., Lee, T.H., Zhang, L., Huang, R.T., et al. (2025). Mechanosensitive Endothelial METTL7A Regulates Internal m (7) G mRNA Methylation and Protects Against Atherosclerosis. bioRxiv. 10.1101/2025.05.22.655328.

57. Piletska, E., Thompson, D., Jones, R., Cruz, A.G., Poblocka, M., Canfarotta, F., Norman, R., Macip, S., Jones, D.J.L., and Piletsky, S. (2022). Snapshot imprinting as a tool for surface mapping and identification of novel biomarkers of senescent cells. Nanoscale Adv 4, 5304–5311. 10.1039/d2na00424k.

58. Kiss, T., Balasubramanian, P., Valcarcel-Ares, M.N., Tarantini, S., Yabluchanskiy, A., Csipo, T., Lipecz, A., Reglodi, D., Zhang, X.A., Bari, F., et al. (2019). Nicotinamide mononucleotide (NMN) treatment attenuates oxidative stress and rescues angiogenic capacity in aged cerebromicrovascular endothelial cells: a potential mechanism for the prevention of vascular cognitive impairment. Geroscience 41, 619–630. 10.1007/s11357-019-00074-2.

59. Wang, L., Mao, B., Fan, K., Sun, R., Zhang, J., Liang, H., and Liu, Y. (2022). ROS attenuates TET2-dependent ZO-1 epigenetic expression in cerebral vascular endothelial cells. Fluids Barriers CNS 19, 73. 10.1186/s12987-022-00370-8.

60. Xiao, P., Zhang, Y., Zeng, Y., Yang, D., Mo, J., Zheng, Z., Wang, J., Zhang, Y., Zhou, Z., Zhong, X., and Yan, W. (2023). Impaired angiogenesis in ageing: the central role of the extracellular matrix. J Transl Med 21, 457. 10.1186/s12967-023-04315-z.

61. Dunn, J., Moore, C., Kim, N.S., Gao, T., Cheng, Z., Jin, P., Ming, G.L., Qian, J., Su, Y., Song, H., and Zhu, H. (2025). Transcription Factor-Wide Association Studies to Identify Functional SNPs in Alzheimer’s Disease. J Neurosci 45. 10.1523/JNEUROSCI.1800-24.2024.

62. Galaxy, C. (2024). The Galaxy platform for accessible, reproducible, and collaborative data analyses: 2024 update. Nucleic Acids Res 52, W83–W94. 10.1093/nar/gkae410.

63. Luo, W., Pant, G., Bhavnasi, Y.K., Blanchard, S.G., Jr., and Brouwer, C. (2017). Pathview Web: user friendly pathway visualization and data integration. Nucleic Acids Res 45, W501–W508. 10.1093/nar/gkx372.

64. Anders, S., Pyl, P.T., and Huber, W. (2015). HTSeq--a Python framework to work with high-throughput sequencing data. Bioinformatics 31, 166–169. 10.1093/bioinformatics/btu638.

65. Love, M.I., Huber, W., and Anders, S. (2014). Moderated estimation of fold change and dispersion for RNA-seq data with DESeq2. Genome Biol 15, 550. 10.1186/s13059-014-0550-8.

66. Dobin, A., Davis, C.A., Schlesinger, F., Drenkow, J., Zaleski, C., Jha, S., Batut, P., Chaisson, M., and Gingeras, T.R. (2013). STAR: ultrafast universal RNA-seq aligner. Bioinformatics 29, 15–21. 10.1093/bioinformatics/bts635.

67. Ewels, P., Magnusson, M., Lundin, S., and Kaller, M. (2016). MultiQC: summarize analysis results for multiple tools and samples in a single report. Bioinformatics 32, 3047–3048. 10.1093/bioinformatics/btw354.

68. Mehta, S., Bernt, M., Chambers, M., Fahrner, M., Foll, M.C., Gruening, B., Horro, C., Johnson, J.E., Loux, V., Rajczewski, A.T., et al. (2023). A Galaxy of informatics resources for MS-based proteomics. Expert Rev Proteomics 20, 251–266. 10.1080/14789450.2023.2265062.

69. Grabherr, M.G., Haas, B.J., Yassour, M., Levin, J.Z., Thompson, D.A., Amit, I., Adiconis, X., Fan, L., Raychowdhury, R., Zeng, Q., et al. (2011). Full-length transcriptome assembly from RNA-Seq data without a reference genome. Nat Biotechnol 29, 644–652. 10.1038/nbt.1883.

